# ChemPrint: An AI-Driven Framework for Enhanced Drug Discovery

**DOI:** 10.1101/2024.03.22.586314

**Authors:** Tyler J. Umansky, Virgil A. Woods, Sean M. Russell, Davey M. Smith, Daniel J. Haders

**Affiliations:** Model Medicines, La Jolla, California, United States; University of California San Diego, La Jolla, California, United States

## Abstract

Traditional High-Throughput Screening (HTS) drug discovery is inefficient. Hit rates for compounds with clinical therapeutic potential are typically 0.5% and only up to 2% maximally. Deep learning models have enriched screening rates to 28%; however, these results include hits with non-therapeutic relevant concentrations, insufficient novelty to their training set, and traverse limited chemical space. This study introduces a novel artificial intelligence (AI)-driven platform, GALILEO, and the Molecular-Geometric Deep Learning (Mol-GDL) model, ChemPrint. This model deploys both t-distributed Stochastic Neighbor Embedding (t-SNE) data splitting to maximize chemical dissimilarity during training and adaptive molecular embeddings to enhance predictive capabilities and navigate uncharted molecular territories. When tested retrospectively, ChemPrint outperformed a panel of five models for the difficult-to-drug oncology targets, AXL and BRD4, achieving an average AUROC score of 0.897 for AXL and 0.876 for BRD4 using the t-SNE split, compared to benchmark model scores ranging from 0.826 to 0.885 for AXL and 0.801 to 0.852 for BRD4. In a zero-shot prospective study, *in vitro* testing demonstrated that 19 of 41 compounds nominated by ChemPrint against AXL and BRD4 demonstrated inhibitory activity at concentrations ≤ 20 µM, a 46% hit rate. The 19 hits reported an average-maximum Tanimoto similarity score of 0.36 relative to their training set and scores of 0.13 (AXL) and 0.10 (BRD4) relative to clinical stage compounds for these targets. Our findings demonstrate that increasing test set difficulty through training and testing ChemPrint on datasets with maximal dissimilarity enhances the predictive capabilities of the model. This results in the discovery of compound libraries at high hit rates with low therapeutic concentrations and high chemical novelty. Taken together, the proposed platform sets a new performance standard.

## 1. INTRODUCTION

### 1.1 Traditional Drug Discovery

Conventional drug discovery and development requires extensive resources and still has low efficiency. In a typical early drug discovery program, the process of screening thousands of compounds with High-Throughput Screening (HTS) yields a mere ∼2% hit rate, maximally^1^. This inefficiency is compounded by the lead optimization process, which requires refining compounds to achieve the desired potency and to address a full Target Product Profile (TPP). In total, 10-15 years^2^ and up to ∼$6 billion in capital are typically required to discover a compound and bring it to FDA approval^3^. This overall inefficiency of traditional drug discovery practices has necessitated the development of new methods, including the use of artificial intelligence (AI).

### 1.2 Challenges with AI Drug Discovery

To date, no AI-discovered therapeutic has achieved FDA approval, and few have progressed beyond preclinical animal model proof of concept^4^. Two reasons for this limited success are that AI platforms often struggle to sufficiently improve prospective discovery accuracy and to identify potent chemistry significantly different from that of known active molecules. The inefficiency of these models is evident in published *in vitro* hit rates from the platforms of venture-backed and public AI drug discovery companies. Based on the hypothesis that commercial platforms can consistently perform well across multiple protein targets and can discover entire libraries of novel hits rather than just isolated successes, we compare the hit rates of AI drug discovery platforms that could be averaged across at least two different targets and where at least ten compounds were screened per target. Schrödinger has reported a pipeline-wide average *in vitro* hit rate of 26% over 14 targets, but also includes hits at concentrations up to 30 μM^5^. Atomwise published an average hit rate of 8.8% across 22 targets; however, this encompasses hits at concentrations up to 250 μM^6^. Insilico Medicine has reported an average of 28% across 2 targets, though this includes hits at concentrations up to 25 μM and lead optimization-phase hits^7,8,9^.

These AI approaches not only have limited hit rates but also report non-standardized concentrations to define “hits”, likely leading to inflated hit rates that permit concentrations that are too high to be therapeutically viable^5,6,7,8,9^. Furthermore, few of the compounds discovered as hits demonstrate significant chemical novelty, highlighting the inability of these models to predict properties of compounds beyond the chemical space they have seen^10,11^. One reason for this, is that standard approaches to model validation inadvertently bias models, limiting their competency to the same chemical space as their training data. These challenges arise because AI models are trained and tuned on datasets lacking chemical diversity and evaluated using conventional performance indicators. While this method shows the models’ technical ability to identify hits from past (retrospective) data, it fails to predict and discover drugs that will have a real clinical impact in future (prospective) studies. New evaluation strategies must be deployed that prioritize the extrapolative ability of AI models to infer beyond known chemical territories.

### 1.3 Model Training Environments

Data splitting methodology for training, validation, and testing sets prior to machine learning (ML) model training and tuning has a significant impact on prediction accuracy. In most ML applications, the common method for splitting data, called ‘stratified random splitting’, is used^13^ (**Figure 1A**). However, this method does not work well for ML drug prediction models because it fails to properly evaluate their performance^13^. Test sets generated by this method of splitting often present models with an oversimplified prediction task. Feinberg et al. have demonstrated that random splitting significantly overestimates the generalization and extrapolation capabilities of ML methods in drug discovery^13^. Alternative dataset splitting techniques, like the temporal split method utilized by Feinberg et al., aim to mitigate this by organizing data based on the publication order^13,14^. This method operates on the principle that the sequential evolution of substructural chemistries over time can provide a more realistic challenge when splitting data for model validation. However, assuming publication dates adhere to medicinal chemistry pedigree can introduce discrepancies, as these dates may not align with the actual order of discovery. While the intention behind the temporal split employed is creative, *in vitro* results that would validate its impact on real-world drug discovery remain unpublished. To this end, we evaluated a novel train-test splitting methodology to directly maximize dataset dissimilarity and challenge the model in an effort to find new chemical novelty (**Figure 1B**).

**Figure 1.**
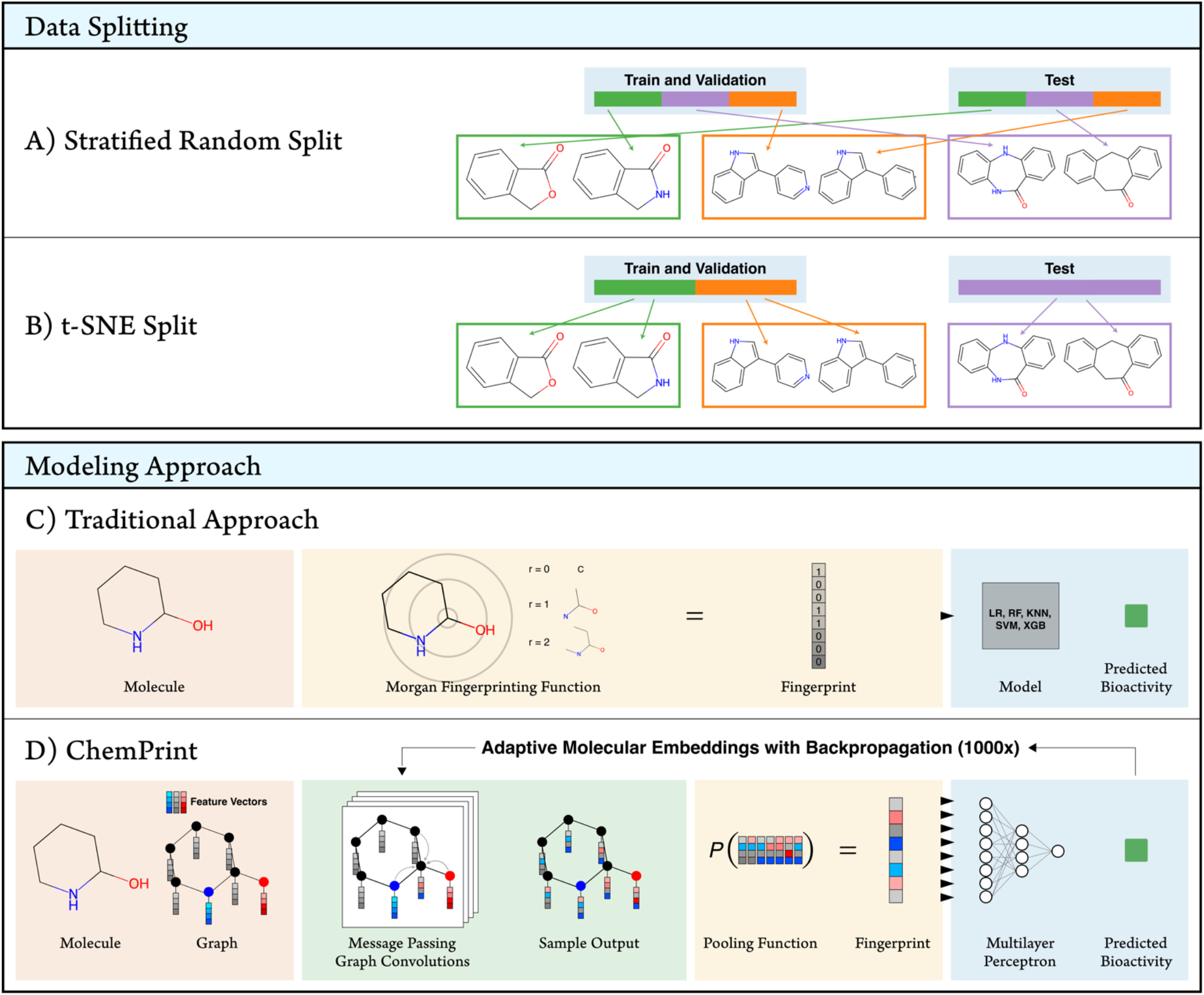
**(A)** Stratified Random Split: In this approach, the dataset is randomly divided into training, validation and test sets. The illustration shows how different groups of molecular structures (green, orange, purple) are distributed across the training set, validation set and test set. This distribution process leads to neighboring pairs of compounds appearing in the training and test set or the validation and test set, for example. This data leakage between data sets maximizes retrospective analysis scores, but limits the models ability to extrapolate to novel chemical space in prospective drug discovery studies. **(B)** t-SNE Split: This approach uses t-Distributed Stochastic Neighbor Embedding (t-SNE) to partition the dataset. Molecules are clustered based on their structural similarity in a reduced-dimensional space. The illustration shows molecules of the same chemical class (green, orange, purple) grouped together within the same dataset. Molecules within the training and validation set are different from those in the unobserved test set. This splitting methodology creates a retrospective training environment where the test set consists of molecules that are structurally different from those in the training and validation sets, which more closely resembles the prospective drug discovery environment for novel discovery and enhances the models ability to extrapolate to novel chemical space. **(C)** Traditional Modeling Approach: This method involves converting the molecule’s structure into a fingerprint using the Morgan fingerprinting function (e.g., ECFP), which encodes structural features into a binary vector. This fingerprint is then used as input for machine learning models such as SVM (Support Vector Machine), RF (Random Forest), LR (Logistic Regression), XGB (Extreme Gradient Boosting), or KNN (K-Nearest Neighbors) to predict the bioactivity of the molecule. As shown, the standard fingerprint is a static embedding that is bounded by a predetermined encoding that is not updated with backpropagation, limiting its competency. **(D)** ChemPrint Approach: This method represents the molecule as a graph, where atoms and bonds are treated as nodes and edges, respectively. Each node is initialized with select feature vectors, and message passing graph convolutions update and refine the feature vectors for each node. These are pooled to form a molecular fingerprint, which serves as input to a multilayer perceptron model to predict bioactivity. The feedforward prediction results backpropagate to redefine model weights 1000 times, creating an adaptive molecular embedding cycle that enhances predictive capabilities and aids the navigation of uncharted molecular territories.

### 1.4 Adaptive Molecular Embeddings

Drug discovery models powered by AI are limited by molecular embeddings that do not evolve or adapt as the model gains knowledge. The reliance on static molecular embeddings, such as Morgan Fingerprints like Extended Connectivity FingerPrints (ECFP),^12^ limits the ability to discover new chemical entities active against the target. Specifically, ECFP encodings use bits that treat similar molecular characteristics as if they were unrelated, thus requiring machine learning algorithms to relearn relationships^13^ (**Figure 1C**). To address this, we used an adaptive molecular embedding technique that evolves with the model’s learning process, enabling the *in silico* identification of multiple novel therapeutic candidates (**Figure 1D**).

### 1.5 Case Study

In this study, we evaluate our AI drug discovery platform (GALILEO) and its zero-shot capable Molecular-Geometric Deep Learning (Mol-GDL)^15^ model (ChemPrint) to target the oncogenic AXL (Gas6/AXL pathway^16^) and BRD4 (BET family^17^) proteins. These two targets were chosen because they affect multiple cancer types, such as breast cancer, pancreatic cancer, and glioblastoma^16,17^ and have demonstrated particularly low hit rates in the post HTS steps of drug discovery, rendering them the classification of ‘undruggable’^18,19^. Supported by *in vitro* experiments, our data indicate significant progress in developing AI that can discover drugs for challenging targets. Importantly, GALILEO was designed to be versatile and can be applied to a wide range of therapeutic areas beyond AXL and BRD4.

## 2. RESULTS

### 2.1 Data Preparation

#### 2.1.1 AXL and BRD4 Data Curation and Binarization

To discover potent, novel compounds that have high drug discovery potential, it is critical that training datasets are built-to-purpose, meaning they are specifically optimized for a particular target. Using our GALILEO data acquisition and curation pipeline, we procured large, high-quality datasets (see methods) from primary source literature for AXL and BRD4. To identify the most highly potent compounds, we defined compounds with a bioactivity measurement below 1 µM as active and above as inactive (**Table 1**).

**Table 1.** Active and Inactive Counts for AXL and BRD4 datasets using a 1 µM threshold.

| Target | Actives | Inactives | Total |
| --- | --- | --- | --- |
| AXL | 1823 | 982 | 2805 |
| BRD4 | 3668 | 1987 | 5655 |

#### 2.1.2 AXL and BRD4 Train-Test Splits

We split the AXL and BRD4 data into training and testing sets based on an 80%-20% stratified random split as well as our proposed splitting methodology that utilized t-distributed Stochastic Neighbor Embedding (t-SNE)^20^ (see methods). This t-SNE splitting protocol aimed to increase the structural and pharmacophoric dissimilarity between training and testing sets, allowing an evaluation of the model’s ability to generalize to new, unseen chemical compounds. Visualization of the t-SNE split revealed clustering of molecules with similar structural and pharmacophoric properties (**Figure 2**). To imitate a data-limited scenario, the cutoff boundary was drawn to create a 50-50% proportion of data within the split train-test sets. Similarly typed compounds were retained in either the train or test set, resulting in two chemically distinct data sets. To demonstrate the chemical dissimilarities between the two datasets, we calculated their average-maximum test-train Tanimoto coefficients^21,22^.

**Figure 2.**
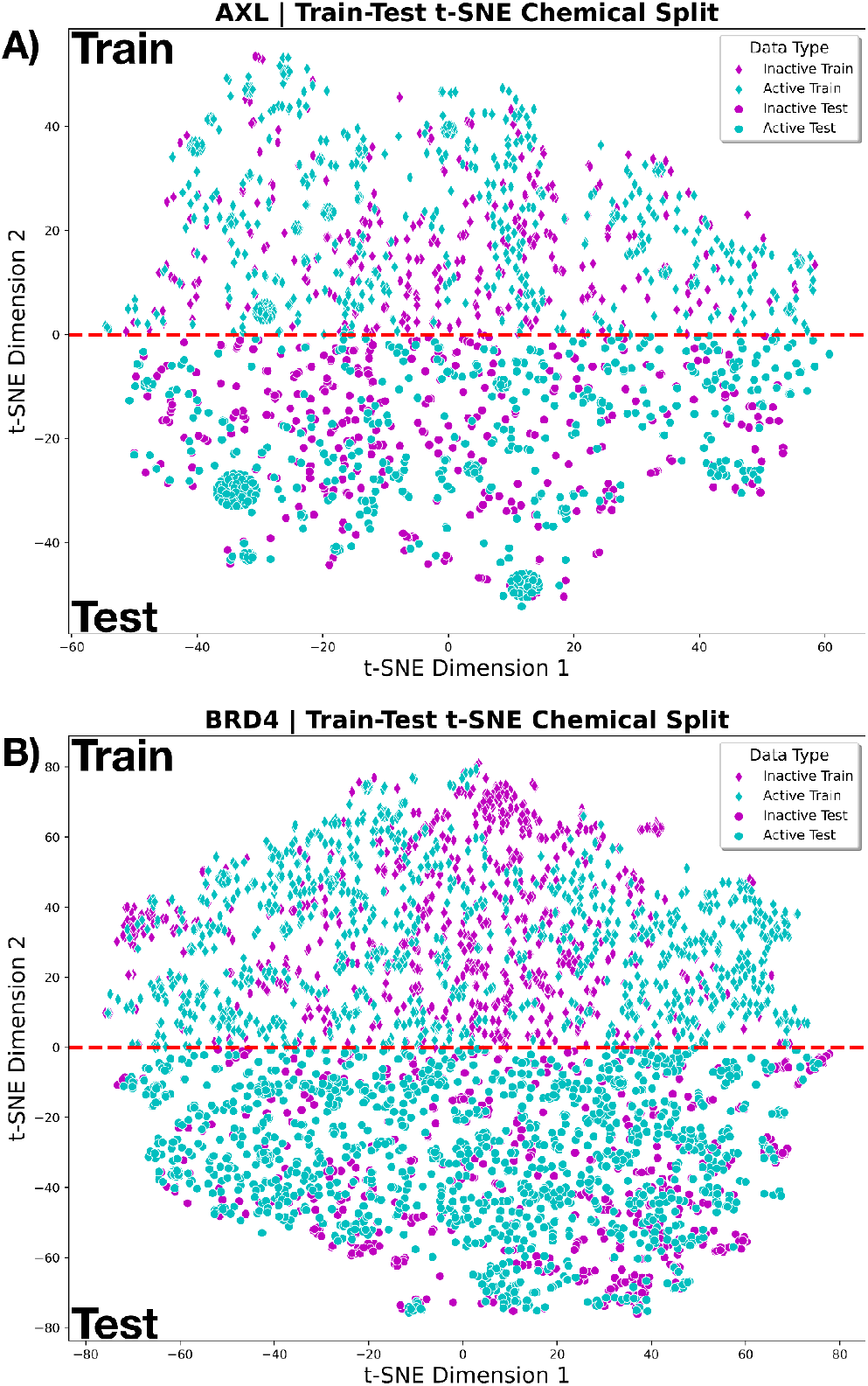
t-SNE visualization of train-test splits for (**A**) AXL and (**B**) BRD4. Points represent compounds: inactive train (magenta diamonds), active train (cyan diamonds), inactive test (magenta circles), and active test (cyan circles). The red line at y = 0 separates the training set (above) from the testing set (below). Clustering patterns indicate distinct chemical environments in each set, demonstrating effective separation.

For the stratified random splits, the average-maximum Tanimoto similarity scores were 0.75 for AXL and 0.80 for BRD4, while for the t-SNE splits, the scores were 0.59 and 0.53, respectively (**Table 2)**. Thus, comparing the splitting methods there was a 27% increase in chemical dissimilarity for AXL and 51% for BRD4 when using the t-SNE method versus the stratified random method. In other words, we observed that the t-SNE method provided better chemical separation between train and test data for both AXL and BRD4. In principle, this t-SNE split added the chemical diversity challenge needed to evaluate a model capable of accurate extrapolation outside of compounds similar to those in the training set.

**Table 2.** Stratified Random and t-SNE Data Split Comparison for AXL and BRD4 datasets.^α^.

| Target | Split Method | Train Actives | Train Inactives | Test Actives | Test Inactives | Tan Similarity |
| --- | --- | --- | --- | --- | --- | --- |
| AXL | SR | 1456 | 785 | 367 | 197 | 0.75 |
|  | t-SNE | 931 | 454 | 892 | 528 | **0.59** |
| BRD4 | SR | 2932 | 1589 | 736 | 398 | 0.80 |
|  | t-SNE | 1639 | 1164 | 2029 | 823 | **0.53** |

<sup>a</sup>SR: Stratified Random Split, Tan: Tanimoto

### 2.2 Performance Evaluation: ChemPrint vs. Benchmark Models

We performed a comparative case study analyzing the performance of our zero-shot, Mol-GDL model, ChemPrint and five benchmark models: logistic regression (LR), random forest (RF), k-nearest neighbors (KNN), support vector machine (SVM), and extreme gradient boosting (XGB)^23,24^. To further validate our t-SNE data splitting method on AXL and BRD4 datasets, we assessed the performance of the models with both our t-SNE split and the stratified random split.

Each model architecture and splitting technique pair was trained and tested ten times, with the Area Under the Receiver Operating Characteristic (AUROC) value recorded for each replicate. A high AUROC means that the model is very good at distinguishing between the different classes (e.g., active vs. inactive compounds). The averages of these AUROC values are reported in **Table 3**, and the single best model run is reported as an ROC curve in **Figure 3**.

**Table 3.** Benchmarking Test Set Performances (Average AUROC ± 95% Confidence Interval)^α^.

|  | AXL |  | BRD4 |  |
| --- | --- | --- | --- | --- |
|  | t-SNE | Stratified Random | t-SNE | Stratified Random |
| ChemPrint | **0.897 ± 0.002** | 0.934 ± 0.001 | **0.876 ± 0.003** | 0.948 ± 0.001 |
| SVM | 0.885 ± 0.000 | 0.949 ± 0.000 | 0.852 ± 0.000 | 0.945 ± 0.000 |
| RF | 0.879 ± 0.003 | **0.961 ± 0.001** | 0.842 ± 0.004 | **0.951 ± 0.001** |
| LR | 0.871 ± 0.000 | 0.957 ± 0.000 | 0.814 ± 0.000 | 0.941 ± 0.000 |
| XGB | 0.872 ± 0.000 | 0.938 ± 0.000 | 0.806 ± 0.000 | 0.929 ± 0.000 |
| KNN | 0.826 ± 0.000 | 0.930 ± 0.000 | 0.801 ± 0.000 | 0.936 ± 0.000 |
| Average of Each Split | 0.872 | 0.945 | 0.832 | 0.942 |

<sup>a</sup>Metric scores for all evaluated models using both the stratified random split and t-SNE split on AXL and BRD4 data. Statistics are averaged over ten distinct model trainings. SVM: Support Vector Machine, RF: Random Forest, LR: Logistic Regression, XGB: Extreme Gradient Boosting, KNN: K-Nearest Neighbors.

**Figure 3.**
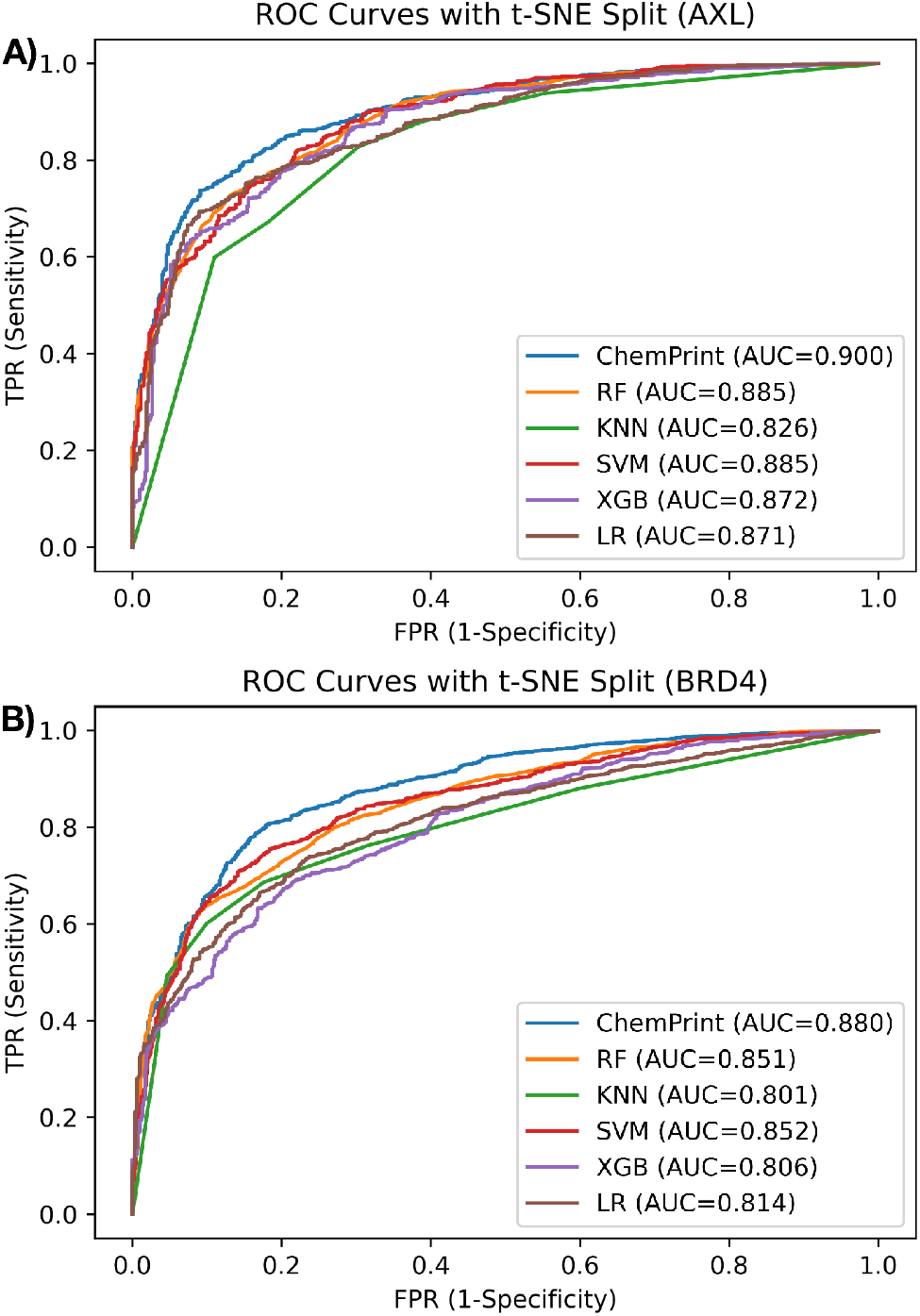
Reported ROC curves and associated AUROC values for the best performing model across all replicates for (**A**) AXL and (**B**) BRD4. Metrics are generated from our t-SNE test split.

For both AXL and BRD4, the ChemPrint models with t-SNE splitting demonstrated the highest performance based on both average (**Table 3**) and single best AUROC scores (**Figure 3**). With the t-SNE split, ChemPrint had an average AUROC score of 0.897 for AXL, whereas other model scores ranged from 0.826 to 0.885. ChemPrint also outperformed all other models for BRD4 when using the t-SNE split with an average AUROC of 0.876 compared to the benchmark models that range from 0.801 to 0.852. The best ChemPrint iterations had higher AUROC scores than each other model across both proteins for the t-SNE split, with score improvements of up to ∼10%, as seen in **Figure 3**.

However, on average, across the six models, the AUROC scores for t-SNE splits were approximately ∼10% lower than those for random stratified splits. This was expected because the t-SNE split created a more challenging evaluation set by increasing the dissimilarity between training and testing sets, better assessing the model’s generalization capability. For AXL models, switching from stratified random splitting to t-SNE splitting resulted in a decrease in mean AUROC from 0.945 to 0.872 (∼8% decrease). Similarly, BRD4 models showed a decrease from 0.942 to 0.832 (∼12% decrease). Examining the average-maximum Tanimoto similarity scores (**Table 2**) in relation to AUROC results, we observed that a decrease in test-train similarity led to a corresponding decrease in AUROC. This suggested that higher test-train similarity (as seen with stratified random splits) inflated AUROC scores, which can be misleading regarding the model’s ability to generalize. In contrast, the t-SNE split created a more rigorous and representative evaluation of the model’s extrapolative ability by increasing the dissimilarity between molecules in the training and test datasets.

### 2.3 Quality Assessment of Adaptive Molecular Embeddings

To elucidate the underlying adaptive molecular embedding process of ChemPrint, we plotted the t-SNE of model latent embeddings at each epoch. The t-SNE visualizations demonstrate that at training initiation (epoch 0), activity classes were largely intersecting embeddings, with no obvious pattern evident (**Figure 4A**), suggesting that no chemical or structural information has yet been learned by ChemPrint (i.e., there was no separation between active and inactive compounds and clear relationships between compounds were not present). As training epochs progressed, the latent representations generated by ChemPrint demonstrated an evolution in embedded learning on molecular structure and activity (**Figure 4B**). Specifically, there was prominent clustering of active and inactive compounds. With few errors in assignments, the active and inactive clusters also served as powerful predictors of activity for unseen compounds in the respective ligand spaces by assigning the label of the nearest cluster to the point in consideration (**Figure 4C-E**).

**Figure 4.**
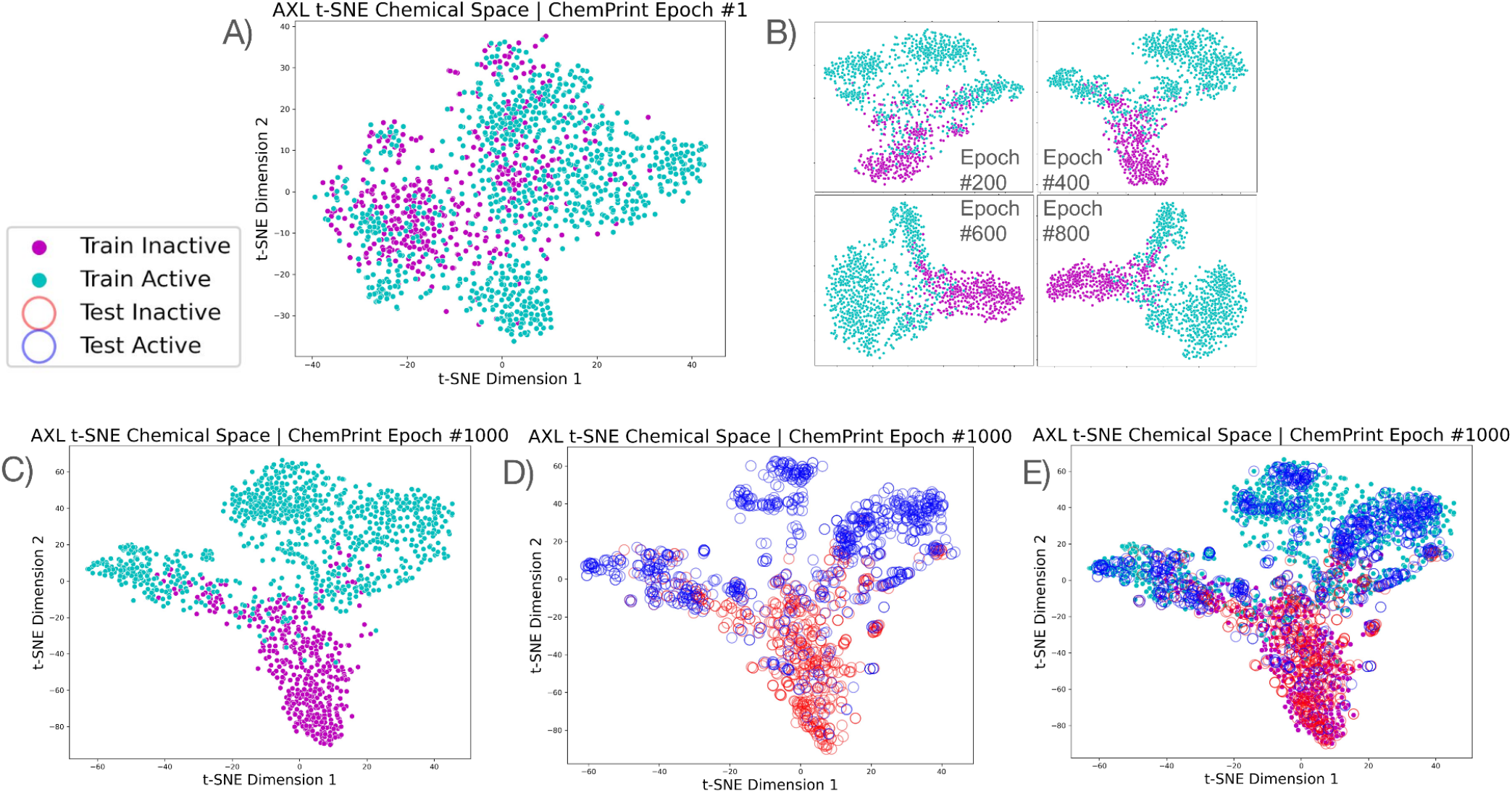
t-SNE visualizations of model latent embeddings of AXL t-SNE split data during ChemPrint’s training cycles. **(A)** t-SNE plot at ChemPrint epoch 1 (initialization). At this stage, active (teal) and inactive (magenta) training compounds are largely overlapping, indicating no learned chemical or structural information. **(B)** t-SNE plots at intermediate training epochs (200, 400, 600, and 800). As training progresses, clustering of active (teal) and inactive (magenta) training compounds begins to emerge, demonstrating the model’s ability to differentiate between active and inactive compounds based on learned molecular features. **(C)** t-SNE plot at ChemPrint epoch 1000 (training). Clear and distinct clusters of active (teal) and inactive (magenta) training compounds are visible, indicating successful embedding of molecular activity information. **(D)** t-SNE plot at ChemPrint epoch 1000 (inference). Here, active (blue) and inactive (red) test compounds are plotted, showing that the model can generalize its learned embeddings to unseen compounds, with test compounds correctly clustering near their respective training clusters. **(E)** Combined t-SNE plot at ChemPrint epoch 1000 (training and inference). This plot includes both training (teal and magenta) and test (blue and red) compounds, demonstrating the overall effectiveness of the embeddings in predicting the activity of unseen compounds based on their proximity to the nearest training clusters.

### 2.4 ChemPrint Inference Predictions

Ensembles of 100 independently trained ChemPrint models for each target were trained on complete curated AXL and BRD4 datasets and deployed to predict activity against compounds in our known chemical entity (KCE) inference library. This library contained approximately 17,000 compounds selected based on having in-human safety and tolerability data, typically from Phase 1 and 2 clinical trials. All compounds from our KCE library with predicted activity against AXL and BRD4 were priority-ranked and then screened by our Freedom To Operate (FTO) pipeline for novelty to target, novelty to indication, and purchasability (see methods). The FTO pipeline identified 41 compounds that met the criteria for novelty to target, novelty to indication, and purchasability. These 41 compounds were subsequently evaluated *in vitro* against their respective targets.

### 2.5 *In Vitro* Compound Validation

The 41 molecules predicted by ChemPrint were tested *in vitro* for enzymatic inhibition via an AXL kinase activity assay and a BRD4 (BD1 + BD2) bromodomain activity assay (see methods). We observed that 19 of the 41 predicted molecules displayed inhibition at ≤ 20 µM, resulting in a 46% hit rate (**Table 4**). The structure of Compound 9 is drawn in **Figure 5A**.

**Table 4.**
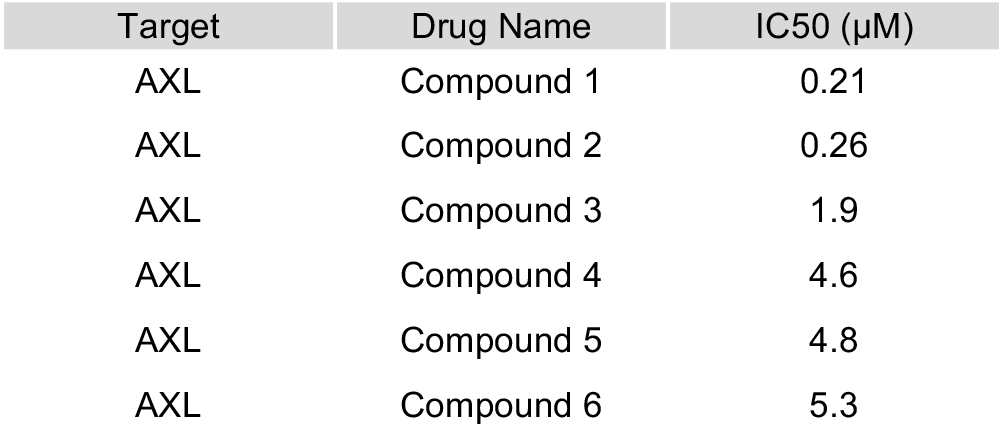
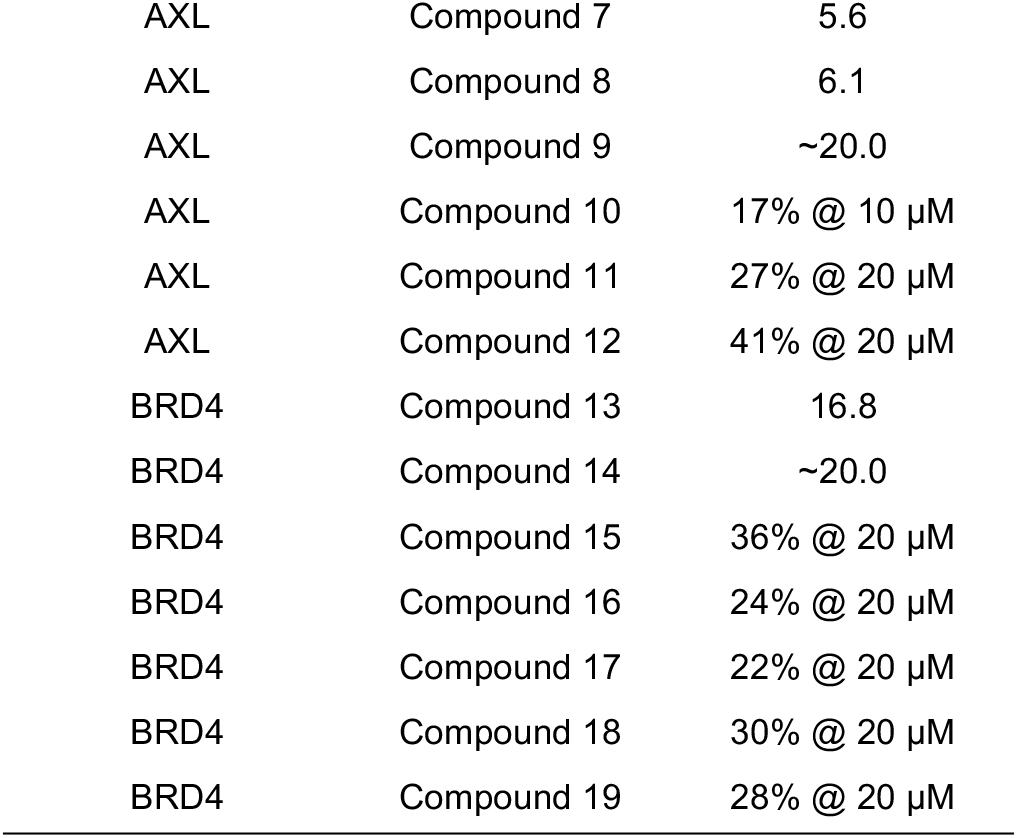
Contains IC50 values of compound inhibitory effects on AXL and BRD4 activities.

**Figure 5.**
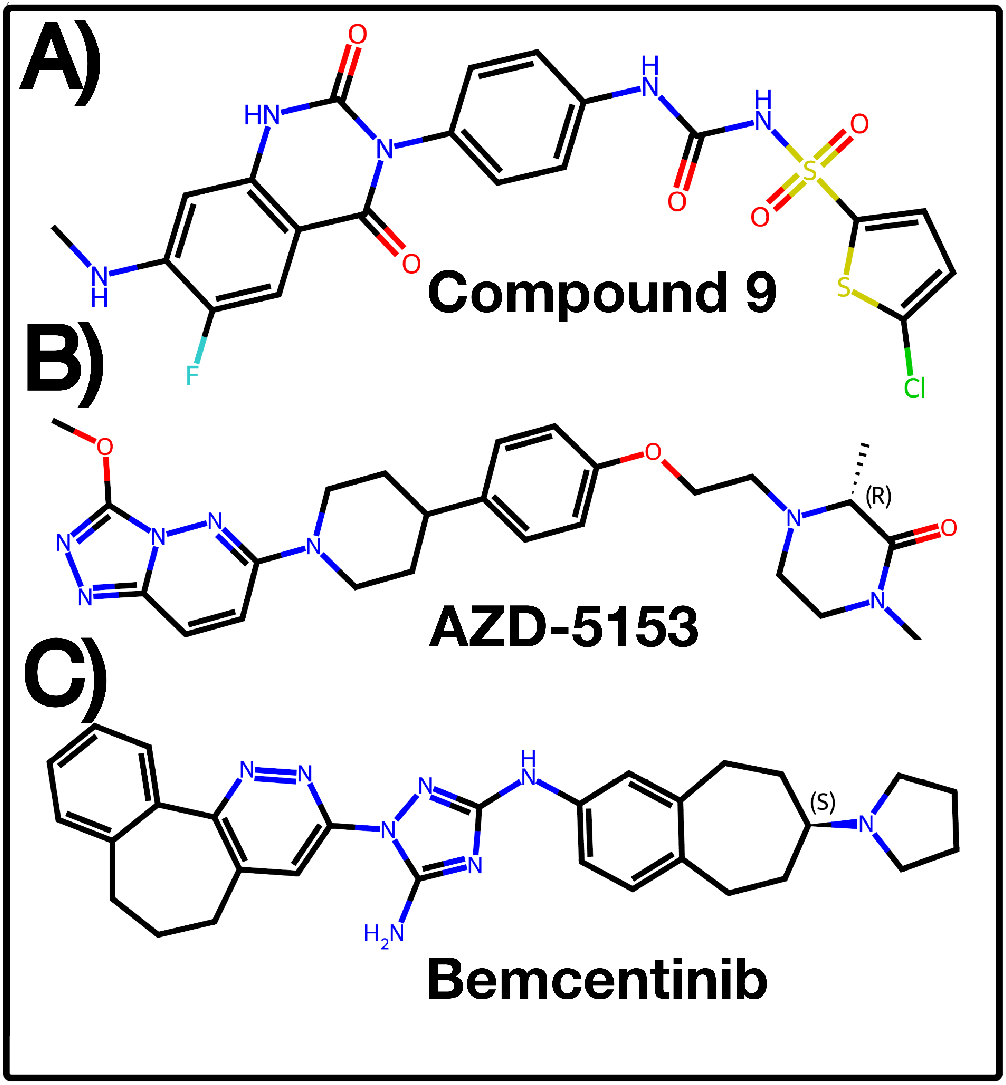
Chemical structures of **(A)** Compound 9, **(B)** AZD-5153, and **(C)** Bemcentinib. ChemPrint discovered Compound 9 as an inhibitor of AXL with an IC50 = 20 µM. AZD-5153 is a potent BRD4 inhibitor actively recruiting for phase 1b/2 clinical trials^25^. Bemcentinib is a potent and selective inhibitor of AXL and has completed phase 2 clinical trials^26^.

### 2.6 Chemical Dissimilarity

To quantify the chemical novelty of our findings, we conducted four types of Tanimoto similarity analyses on our 19 validated hits using ECFP4 fingerprints. The analyses were performed against the training data used for their discovery, post-Phase 1a AXL and BRD4 clinical trial compounds (AZD-5153 and Bemcentinib) (**Figure 5B-C**), all available ChEMBL data for each target, and a pairwise comparison among the hits themselves. First, we recorded the maximum Tanimoto similarity between each *in vitro* hit and both active and inactive compounds from the training set. The similarity means were 0.40 for AXL and 0.30 for BRD4, with an average of 0.36 across both targets (**Figure 6A**). Second, we recorded the maximum Tanimoto similarity between each *in vitro* hit and all ChEMBL compounds for each target. The mean similarity across both targets was 0.37 (**Figure S1**). Third, we recorded the Tanimoto similarity between each *in vitro* hit and Bemcentinib for AXL and AZD-5153 for BRD4. The similarity means were 0.13 for AXL and 0.10 for BRD4 (**Figure 6B**). Fourth, we calculated the mean pairwise Tanimoto similarity among the hits themselves. The mean pairwise similarity was 0.17 for AXL and 0.11 for BRD4, with an average of 0.14 across both targets. These scores indicate that our hits are chemically novel, possessing distinct properties compared to known inhibitors of AXL and BRD4, and the pairwise similarity analysis further demonstrates the diversity within our set of hits.

**Figure 6.**
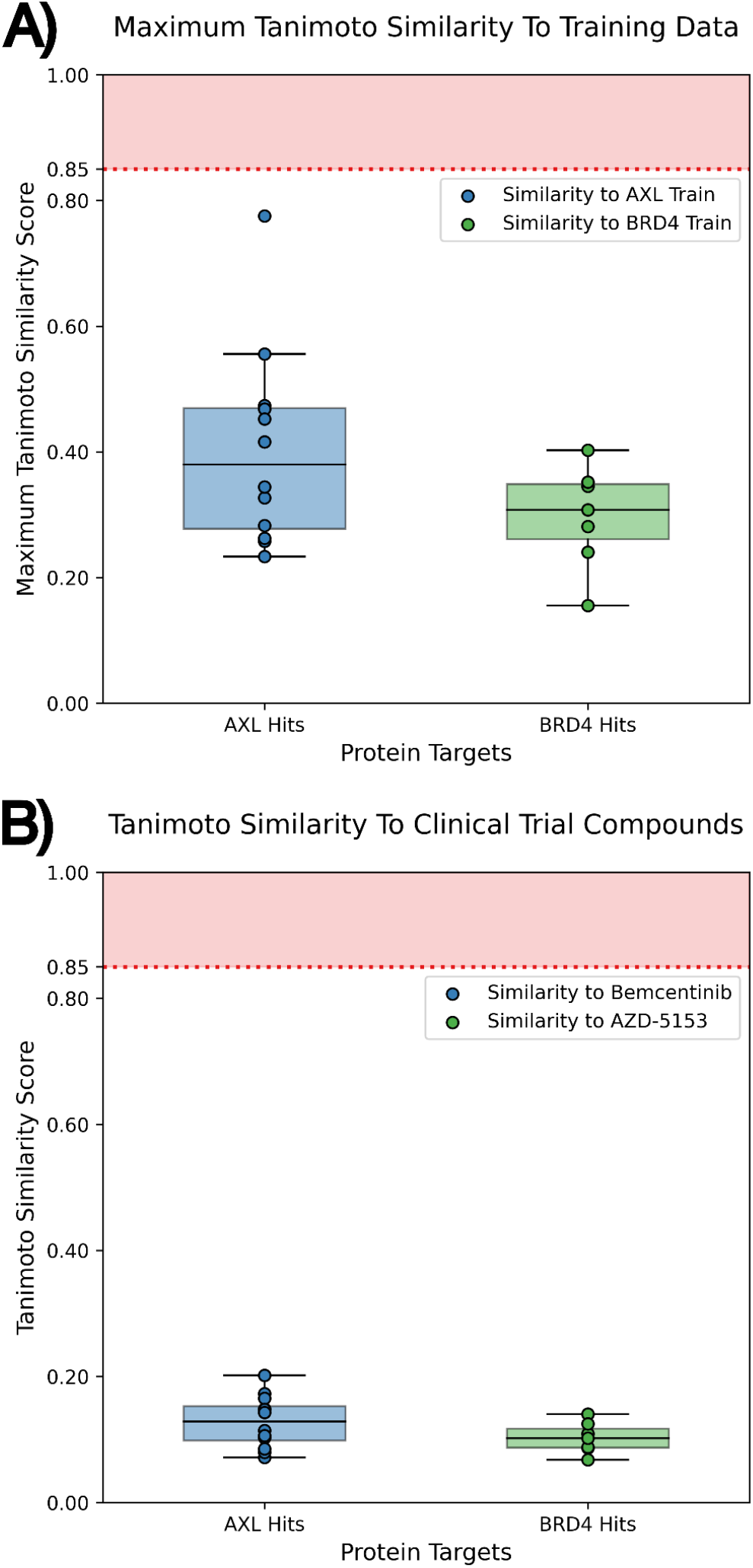
Box and scatter plots of our *in vitro* validated hits for both AXL and BRD4, displaying their **(A)** maximum Tanimoto similarity scores to the training datasets used for their discovery and **(B)** Tanimoto similarity scores to Bemcentinib and AZD-5153, respectively.

## 3. DISCUSSION

### 3.1 Advancements in AI-Driven Drug Discovery

This study had four main findings. First, we found significant improvement in chemical dissimilarity between training and testing datasets when using t-SNE compared to stratified random splitting. Such diversity of chemical entities in datasets improves the ability of AI-driven models to find chemically and mathematically novel active chemical compounds against the target. Second, we found that the AI-driven ChemPrint model outperformed benchmark models in correctly predicting the confusion matrix on the most difficult test sets in retrospective studies. Third, we found that of the 41 compounds ChemPrint nominated as active against AXL and BRD4, 19 had activity at low concentrations (≤ 20 µM). This 46% hit rate (**Table 4**) for AXL and BRD4 inhibitors is a 64% increase over average rates reported by industry leaders^5,6,7,8,9^. Fourth, by evaluating ChemPrint during its learning process, we could observe the progressive formation of active and inactive clusters that accurately predicted the activity of unknown compounds in their respective ligand spaces.

### 3.2 Data Curation and Splitting

The GALILEO pipeline provided high-quality datasets for robust model training. Defining compounds by bioactivity thresholds helped ensure that the datasets were large and highly specific to the targets of interest (AXL and BRD4), which is essential for finding potent inhibitors. Next, we wanted to diversify our train and test sets since such diversity is needed to find new chemical entities. We found that creating chemically diverse training and testing sets was greatly improved by the use of our newly developed t-SNE splitting method, i.e., lower average-maximum Tanimoto similarity scores than the stratified random split.

### 3.3 Model Performance and Evaluation

We evaluated ChemPrint versus five benchmark models of drug discovery, and we found that AUROC scores for each method were dependent upon splitting technique. As seen in **Table 3**, if the “best” performing model were selected through metric evaluation on the stratified random split, RF would be nominated as the best model, but as demonstrated by the t-SNE split, both ChemPrint and SVM perform better, showcasing their ability to generalize beyond their training data. The stratified random split confounded model performance and failed to provide granular differentiation between model abilities, especially for unseen chemical space. The t-SNE split allows for a better evaluation of models that can prospectively discover active compounds from novel chemical space. ChemPrint’s success is likely due to its Mol-GDL architecture, which uses adaptive molecular embeddings instead of the static fingerprints used by conventional models, thereby enhancing its extrapolative potential.

### 3.4 Adaptive Molecular Embeddings

Adaptive molecular embeddings allowed for continuous learning and refinement, as shown by the clear separations of active and inactive compounds in the visualizations of ChemPrint’s learned embeddings across epochs (**Figure 3**). Such insights should allow human drug developers to understand model performance that moves beyond AUROC and similar metrics. This is a quality that is unique to ChemPrint versus the comparative models, as they learn from static embeddings. This quality of ChemPrint allows for Human-In-The-Loop (HITL) interpretation and Informed Inductive Bias (IIB) and removes reliability concerns associated with alternative black-box AI drug discovery approaches.

### 3.5 Novel Compound Identification and *In Vitro* Validation

ChemPrint screened a KCE inference library of approximately 17,000 compounds and identified 41 molecules as potential hits for AXL and BRD4, which were then tested *in vitro*. Hits were conservatively defined as molecules showing *in vitro* enzymatic inhibition in an AXL kinase activity assay and a BRD4 (BD1 + BD2) bromodomain activity assay at concentrations at/below 20 µM. This resulted in a hit rate of 46% (**Table 4**), which is a 64% increase over the average hit rates reported by industry leaders^5,6,7,8,9^. Importantly, all observed activity occurred at low concentrations (≤ 20 µM). In comparison, hit rates reported by other AI drug discovery companies are generally more permissive. Schrödinger has reported an average *in vitro* hit rate of 26% over 14 targets, but this includes hits at concentrations up to 30 μM and lacks detailed data for a fine-grain analysis of compound diversity^5^. Atomwise published an average hit rate of 8.8% across 22 targets, which encompasses hits at concentrations up to 250 μM and includes compounds that may exhibit binding and not activity^6^. Insilico Medicine reported an average hit rate of 28% across two targets, with hits at concentrations up to 25 μM, including both lead optimization-phase compounds and binders without confirmed activity^7,8,9^. These comparisons highlight the rigorous criteria and effectiveness of our platform in identifying active compounds at lower, therapeutically relevant concentrations.

### 3.6 Chemical Dissimilarity

The similarity analysis of the 19 validated hits revealed considerable chemical novelty, with an average-maximum Tanimoto similarity score of 0.36 to their training set and scores of 0.13 (AXL) and 0.10 (BRD4) relative to clinical stage compounds. In the industry, a Tanimoto similarity score of 0.85 is commonly used as a threshold for chemical similarity^27,28^. Our hits, with scores well below this threshold, demonstrate novel and distinct chemistry. In fact, we set our bar much higher, requiring our average Tanimoto similarity to be less than 0.5. A simple Tanimoto similarity search would have failed to identify these hits, highlighting ChemPrint’s ability to generalize beyond its training data. This capability is particularly important as predicting the efficacy of molecules that do not closely resemble any previously tested compounds remains a significant challenge in computer-aided drug design^11,29^. Additionally, the mean pairwise similarity among the hits themselves was 0.14 across both targets, demonstrating diversity within our set of hits and highlighting the hit-finding capabilities of ChemPrint. ChemPrint’s success in discovering novel and diverse chemistry, dissimilar to previously explored compounds, is likely due to its adaptive fingerprints and our evaluations using the t-SNE split.

### 3.7 Limitations and Future Work

While this study showcases the potential of AI-driven models for drug discovery, limitations remain. First, the reliance on t-SNE splitting may not capture all aspects of chemical diversity, and model performance may vary depending on data quality and diversity. Second, the GALILEO and ChemPrint platforms will likely perform differently for different targets and when exploring different datasets. Third, AI models will continue to improve, which should enhance these and other models. Fourth, the hit rate of 46% for two notoriously difficult targets to drug leads the industry^5,6,7,8,9^, but the found chemical entities have not been tested *in vivo*.

## 4. CONCLUSION

Our technology stack (GALILEO with prospective data segmentation and ChemPrint with adaptive molecular embeddings) nominated 41 new compounds to have a desired therapeutic activity for AXL or BRD4, and 19 of them had such activity at low concentrations with high chemical novelty. Such high hit rates with diverse chemical compounds suggest that the proposed platform may overcome traditional drug discovery barriers and efficiently discover novel therapeutic agents. Furthermore, GALILEO was designed to be versatile and indication-agnostic, offering broad applicability across disease targets beyond AXL and BRD4.

## 5. METHODS

### 5.1 Data Sets and Preparation for Prospective Discovery

#### 5.1.1 Data Curation

Our GALILEO drug discovery pipeline was used to curate protein-ligand interaction data for AXL and BRD4. GALILEO identified primary source literature (**supporting documents**), extracted protein-specific structure-activity relationship data that was checked by our experts and filtered for undesired characteristics, such as Pan Assay Interference Compounds (PAINS)^30^, molecular weight extrema, and duplicate compounds. Based on the distribution of activity values in the training sets, a potency cutoff of 1 µM was applied to separate active from inactive compounds in each set. This cutoff created a balance between active and inactive compounds and minimized data exclusion.

#### 5.1.2 Train-Test Split

To split our data into training and test sets, we utilized t-SNE to project ECFP4 encoded molecule data into 2-dimensions. We used the line y = 0 to separate training (above) and test (below) data. This decision boundary created chemically distinct training data and testing data, while also creating our desired ∼50-50% (within 2%) train-test split that maintains class balance. Code Availability: GitHub repository for the t-SNE splitting method is archived at https://github.com/Model-Medicines/tSNE-Chemical-Data-Splitting.

#### 5.1.3 Split Evaluation

The Tanimoto coefficient^37^ is a common metric to evaluate the chemical similarity between molecules (represented as bit-vectors), and is defined as:

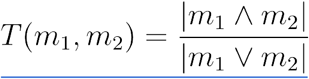

Where |m_1_ ⋀ m_2_ | is the number of common 1’s in the bit-vectors and |m_1_ ⋁ m_2_ | is the number of indices where either bit-vector has a 1. This value ranges from 0 to 1, with a value of 0 indicating completely distinct representations and 1 indicating identical representations. Using this metric, we can approximate the chemical diversity of the dataset split via the average of the maximum Tanimoto coefficient of each test molecule when measured against all training points, i.e.:

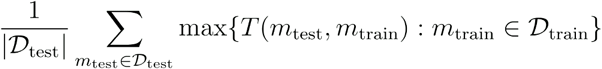

### 5.2 ChemPrint Architecture

ChemPrint leverages Mol-GDL to learn the adaptive embeddings derived from its training data to make predictions when given novel inference sets (**Figure 1D**). The input data takes the form of a geometric graph, a molecular representation that encapsulates the structural information of each datapoint, to which one-hot encoded features are passed in. This graph-based learning enabled us to encode and leverage the inherent dependencies and relationships among the data. ChemPrint architecture encompasses an end-to-end Graph Convolutional Network (GCN) with a Multilayer Perceptron (MLP) module to facilitate positive and negative classification. Select normalization, activation, pooling, and dropout layers are used.

### 5.3 Benchmark Models

ChemPrint was benchmarked against five widely utilized machine learning algorithms. For a molecule to be evaluated by a classifier, it must first be processed into a vectorized numerical representation or embedding that preserved as much structural and chemical information as possible. We utilized the general hashed radial fingerprint, ECFP4 (radius of 2) encoded in 2048 bits to preprocess the data used to train and test the benchmark models. Logistic regression (LR), random forest (RF), k-nearest neighbors (KNN), and support vector machine (SVM) were implemented using the scikit-learn library^23^. The extreme gradient boosting (XGB) was built using the XGBoost library^24^.

### 5.4 Performance Evaluation

#### 5.4.1 Area Under the Receiver Operating Characteristics (AUROC)

The Receiver Operating Characteristic (ROC) curve plots a classifier’s true positive rate (sensitivity) versus its false positive rate (1-specificity) as the classification threshold is varied from 0 to 1. The AUROC curve established relationships between the model’s predicted probabilities of activity and true binding status, reflecting the likelihood of the classifier assigning a higher probability of a positive example (active molecule) compared to a negative example (inactive molecule). AUROC was calculated using the scikit-learn library^23^ for each replicate (n=10) of every model-splitting method pair.

#### 5.4.2 Molecular Embeddings

ChemPrint learns optimal, real-valued, differentiable, property-specific, molecular embeddings by leveraging its GCN instead of the non adaptive ECFP fingerprinting method (used by the LR, RF, KNN, SVM, XGB models). GCN latent space molecular embeddings are engineered and optimized through end-to-end back propagation. In comparison to embeddings such as ECFP, ChemPrint’s embeddings are adaptive and self optimizing for a specific molecular property. **Figure 1D** depicts the graphical representation of the adaptive embedding used as input for our model as compared to the stagnant molecular fingerprint leveraged by benchmark models.

For our analysis of ChemPrint’s embedding space, we captured the model’s learned embeddings at progressive epochs during the training process (**Figure 4**). We then projected these embeddings down to 2 dimensions using t-SNE, and plotted all points colored according to activity and distinguished them as train or test molecules.

### 5.5 ChemPrint Compound Nomination

#### 5.5.1 Known Chemical Entity (KCE) Library

Our known chemical entity (KCE) inference set contains ∼17,000 curated compounds that have been thoroughly characterized and have a documented history in clinical trials. For each compound in this inference set, pertinent information such as synonyms/identifiers, absorption, distribution, metabolism, excretion, toxicity, clinical trial details, current approval statuses, and purchasability are recorded.

#### 5.5.2 Freedom to Operate Pipeline (FTO)

As part of the GALILEO platform of AI drug discovery tools, each predicted active compound from our *in silico* discovery pipeline is investigated for FTO using data miners and web search tools. Non-obviousness and novelty of the predicted compounds are verified using these tools. The data miners make use of our exhaustive set of synonyms and identifiers for each compound to perform in depth target and indication searches. These searches are performed across the six databases ACM, Arxiv, Biorxiv, Medrxiv, Pubmed, and Scopus. Primary source material is returned when both the synonym/identifier and indication or synonym/identifier and target is found in the same publication. The identified literature is then subjected to our natural language processing (NLP) and large language model (LLM) packages to ensure that followup manual review relies on proper relevance and filtration of inconsequential co-mentions of compound and target in the literature.

#### 5.6.1 Enzyme Inhibition Assays

All *in vitro* experiments were conducted by BPS Bioscience Inc.

#### 5.6.1 AXL Kinase Assay

The assay was performed using ADP-Glo Kinase assay reagents (Promega). It measures kinase activity by quantitating the ADP amount produced from the enzymatic reaction. The luminescent signal from the assay is correlated with the amount of ADP present and is directly correlated with the amount of kinase activity. The compounds were diluted in 10% DMSO and 2.5 µl of the dilution was added to a 25 µl reaction so that the final concentration of DMSO is 1% in all of reactions. All of the enzymatic reactions were conducted at 30 ºC for 45 minutes. The 25 µl reaction mixture contains 40 mM Tris, pH 7.4, 10 mM MgCl2, 0.1 mg/ml BSA, 1 mM DTT, 10 µM ATP, kinase substrate and the enzyme. After the enzymatic reaction, 25 µl of ADP-Glo reagent was added and incubated for 45 - 60 minutes at room temperature, followed by another 45 mins incubation with 50 µl of kinase detection mixture. Luminescence signal was measured using a BioTek Synergy 2 microplate reader.

#### 5.6.2 BRD4 (BD1 + BD2) Bromodomain Assay

The assay was performed by TR-FRET technology using recombinant bromodomains (BD1 and BD2) and the endogenous BET Ligand. The TR-FRET signal from the assay is correlated with the amount of Ligand binding to the bromodomain. The compounds were diluted in 100% DMSO. They were diluted 20-fold in 5% DMSO in Reaction Buffer and 2 µl of the dilution was added to a 20 µl reaction so that the final concentration of DMSO is 0.5% in all of reactions. The binding reaction was conducted at room temperature. The 20 µl reaction mixture in Assay Buffer contains the protein, the indicated amount of the inhibitor, acetylated BET ligand (or non-acetylated ligand for negative control wells), and the reaction dyes. The reaction mixture incubated for 120 min prior to reading the TR-FRET signal. Fluorescence signals for both the donor and acceptor dyes were measured using a Tecan Infinite at an excitation of 340 nm and emissions at 620 nm and 665 nm.

## ASSOCIATED CONTENT

## Supporting Information

The Supporting Information is available free of charge on the

ACS Publications website.

Source literature of AXL and BRD4 modeling data (XLSX)

## AUTHOR INFORMATION

## Notes

The authors declare the following competing financial interest(s): D.M.S. receives consulting fees from Bayer, Lucira, Pharma Holdings, Evidera, Vx Biosciences, Gilead, Hyundae, and Red Queen Biosciences, stock options from Fluxergy, Linear Therapies, Vx Biosciences, and Model Medicines, and institutional payments to UCSD from the NIH.

Safety statement: no unexpected or unusually high safety hazards were encountered in this study.

## ACKNOWLEDGMENTS

The authors would like to acknowledge the support and valuable discussions from numerous consultants, advisors, partners, and former employees. Special thanks are due to Dr. David Garvey and Dr. Tushar Menon for their consultation and thorough review of this manuscript and to Navya Ramesh for supporting the team in the construction of benchmark models. The authors acknowledge the contributions of ChatGPT-4 in refining this manuscript’s language to improve coherence and fluency^31^.

## Notes

https://github.com/Model-Medicines/tSNE-Chemical-Data-Splitting.git

### Summary of Updates

The resolution of figures was increased.

