## Supplemental Figures for "ChemPrint: An AI-Driven Framework for Enhanced Drug Discovery"

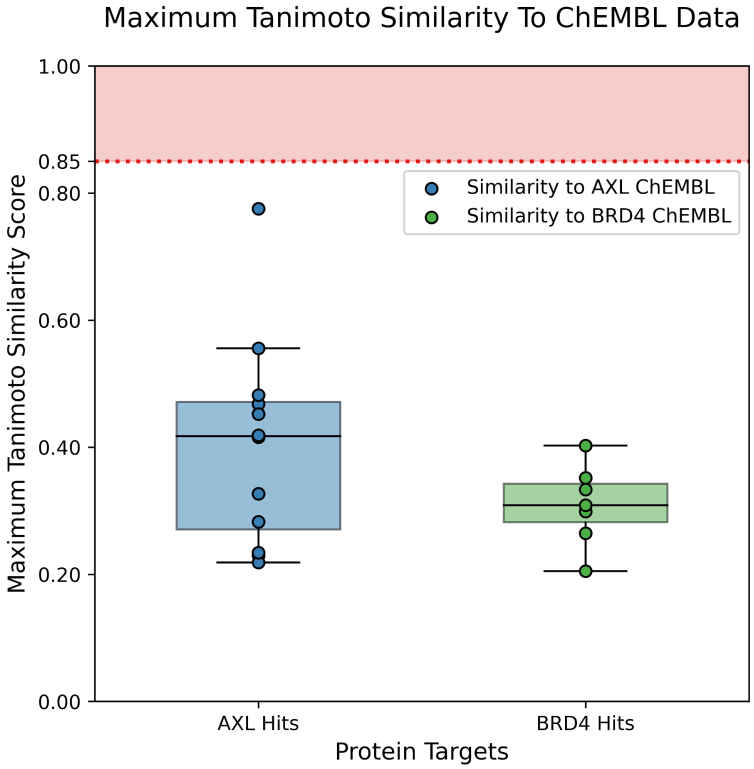


**Figure S1.** Box and scatter plots of our *in vitro* validated hits for both AXL and BRD4, displaying their maximum Tanimoto similarity scores to all available ChEMBL data for each target.

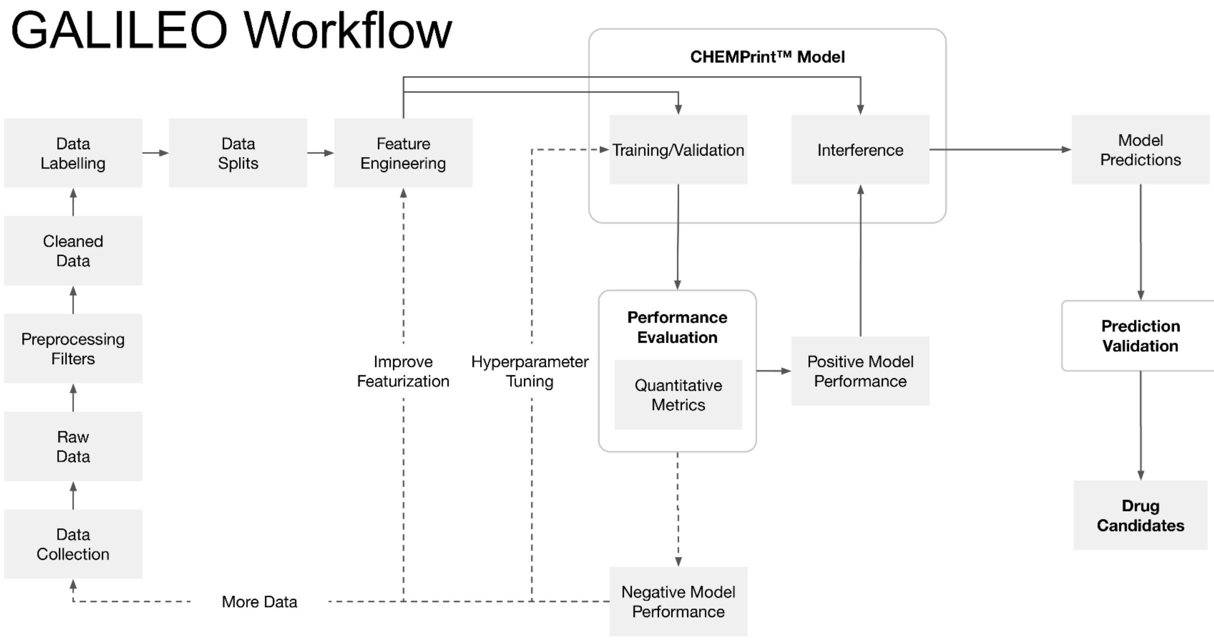


**Figure S2.** Workflow of ChemPrint within our GALILEO AI drug discovery pipeline as detailed in Methods.
